# Estimation of crossbridge-state during cardiomyocyte beating using second harmonic generation

**DOI:** 10.1101/2022.10.13.512034

**Authors:** Hideaki Fujita, Junichi Kaneshiro, Maki Takeda, Kensuke Sasaki, Rikako Yamamoto, Daiki Umetsu, Erina Kuranaga, Shuichiro Higo, Takumi Kondo, Yasuhiro Asano, Yasushi Sakata, Shigeru Miyagawa, Tomonobu M Watanabe

**Affiliations:** Department of Stem Cell Biology, Research Institute for Radiation Biology and Medicine, Hiroshima University, 1-2-3 Kasumi, Minami-ku, Hiroshima, 734-8553, Japan; Laboratory for Comprehensive Bioimaging, RIKEN Center for Biosystems Dynamics Research (BDR), 2-2-3, Minatojima-minamimachi, Chuo-ku, Kobe, 650-0047, Japan; Department of Cardiovascular Surgery, Graduate School of Medicine, Osaka University, 2-2 Yamadaoka, Suita, Osaka 565-0871, Japan; Department of Medical Therapeutics for Heart Failure, Graduate School of Medicine, Osaka University, 2-2 Yamadaoka, Suita, Osaka 565-0871, Japan; Department of Cardiovascular Medicine, Graduate School of Medicine, Osaka University, 2-2 Yamadaoka, Suita, Osaka 565-0871, Japan; Laboratory for Histgenetic Dynamics, Graduate School of Life Sciences, Tohoku University, 6-3 Aramaki Aza-aoba, Aoba-ku, Sendai, Miyagi, 980-8578, Japan

**Keywords:** Cardiomyopathy, Myosin, Second harmonic generation, MYBPC3 deficiency, Radiation effect

## Abstract

Estimation of dynamic change of crossbridge formation in living cardiomyocytes is expected to provide crucial information for elucidating cardiomyopathy mechanisms, efficacy of an intervention, and other parameters. Here, we developed an assay system to dynamically measure second harmonic generation (SHG) polarization in pulsating cardiomyocyte and proved that the SHG anisotropy derived from myosin filaments in disease-model cardiomyocytes depended on their crossbridge status, providing an evaluation method for myosin force generation. Experiments utilizing an inheritable mutation that induces excessive myosin-actin interactions revealed that the correlation between sarcomere length and SHG anisotropy represents crossbridge formation ratio during pulsation. Furthermore, the present method found that ultraviolet irradiation induced an increased population of attached crossbridges that lost force-generating ability upon myocardial differentiation, causing acquired dysfunction. Taking an advantage of infrared two-photon excitation in SHG microscopy, myocardial dysfunction could be intravitally evaluated in a *Drosophila* disease model. Thus, along with the establishment of the methodology, we successfully demonstrated the applicability and effectiveness of the present method to evaluate the actomyosin activity of a drug or genetic defect on living cardiomyocytes.

## Introduction

Cardiomyopathy, a disease of cardiac dysfunction, is the worldwide leading cause of sudden death in young people, including children (Rizzo, Carturan et al., 2019). Muscle fiber contractions in the heart are synchronized through cell adhesion and electrical transmission during normal beating. Abnormalities in any of the elements of an integrated heart system may result in heart failure (Caforio, Pankuweit et al., 2013, Richardson, McKenna et al., 1996). Among them, muscle contraction, the driving force of the heartbeat, is a promising target for understanding the occurrence mechanism and treatment of systolic heart failure (Moore, Leinwand et al., 2012, Walsh, Rutland et al., 2010). In cardiomyocytes, cardiac myosin, a cytoskeletal motor protein, generates contractile forces along an actin filament using energy stored in adenosine triphosphate (ATP) under the control of various regulatory proteins (Barrick & Greenberg, 2021, Hanft, Korte et al., 2008). Muscle force generation dysfunction in heart is mainly caused not by mutation in myosin motor itself but mutations in myosin regulatory/accessory proteins via malfunctions of various pathways including signal transduction, calcium (Ca^2+^) cycling, adenine nucleotides transportation, reactive oxygen species (ROS) production, and so on (Caforio et al., 2013, Richardson et al., 1996). To reveal the causal relationship from the mutation to heart failure, and to investigate the toxicity, the severity, or the efficacy of a drug against cardiomyocytes, it is necessary to quantify their effects on actomyosin activity during cardiac muscle contraction in living cardiomyocytes.

Heartbeat measurement based on video analysis is a simple non-invasive method for evaluating cardiomyocyte dysfunction (Maddah, Heidmann et al., 2015, Santoso, Farhan et al., 2020). However, it is not applicable for selectively focusing on myosin motor function. Changes in the length of the sarcomere, the smallest functional unit of the muscle, is an optically-visible dynamic indicator of muscle function, but is still not sufficient to evaluate actomyosin activity. It would be ideal to directly measure the myosin force generation in living cardiomyocytes. However, various techniques for measuring the forces exerted by myosin molecules, myofibers, and myocytes, developed from a long history of muscle mechanobiology, are almost all contact measurements, such as atomic force microscopy, traction microscopy, and magnet/laser trapping, or invasive measurements, such as laser ablation (Roca-Cusachs, Conte et al., 2017, Woody, Greenberg et al., 2018). This is because force is defined through the properties of a material and its physical deformation. A non-invasive/non-contact method for measuring actomyosin activity in living muscle cells, even without directly measuring force itself, could accelerate elucidation of the mechanism and drug discovery for cardiomyopathy and the research in mechanobiology. In this study, we focused on the structural dynamics of the actomyosin complex during force generation. The structural state of actomyosin in a sarcomere determines the anisotropic features of optical second harmonic generation (SHG) (Forderer, Georgiev et al., 2016, Nucciotti, Stringari et al., 2010, Plotnikov, Millard et al., 2006, Psilodimitrakopoulos, Loza-Alvarez et al., 2014, Schurmann, von Wegner et al., 2010, Yuan, Wang et al., 2019). This indicates that SHG polarization microscopy is a potential alternative to direct force measurements in muscles.

SHG is a nonlinear coherent scattering process that reflects permanent electric dipole moments and their alignments in illuminated materials (James & Campagnola, 2021). The SHG light is strongly generated from fibrous materials with asymmetries in electric polarization, such as fibrillar collagen in tendons, myosin thick filaments in myosarcoma, and densely bundled microtubules (Cicchi, Vogler et al., 2013, Cox, 2011, Mohler, Millard et al., 2003). The coefficients representing SHG, a 3rd-rank polar tensor, originate from the electric dipole moments and can be measured using the polarization dependence of SHG (James & Campagnola, 2021). While both myosin and actin in the actomyosin complex emit SHG, the SHG from myosin is over three orders of magnitude larger than that of actin (Plotnikov et al., 2006). Myosin is composed of a tadpole-shaped subdomain-1 (S1), an α-helical coiled-coil subfragment-2 (S2), and a light meromyosin (LMM) region for rod filamentation (Fig. 1, *left*). The S1 region converts chemical energy from ATP hydrolysis into mechanical energy by changing its structure corresponding to the nucleotide state, called “lever arm swinging.” The LMM is responsible for rod filamentation, while the S2 region acts as a “spring” between the converter and the rod (Barrick & Greenberg, 2021). The primary source of SHG in muscle is thought to be the S2 and the LMM, whose structures comprise highly regular unidirectional filaments (Plotnikov et al., 2006). In contrast, the contribution of the globular S1 to SHG is limited to approximately 10% (Nucciotti et al., 2010). During muscle contraction, S1 is dynamic, LMM is static, and S2 interlocks with S1 dynamics. Therefore, the tilting or bending of the S2 region coupled with myosin force generation alters SHG anisotropy in a contracting sarcomere, which can be detected through the incident polarization dependence of SHG intensity (Fig. 1A). Several groups have reported their success in discriminating SHG anisotropy between rigor and relaxed states in rabbit psoas fibers (Nucciotti et al., 2010), isolated scallop myofibrils (Plotnikov et al., 2006), mouse tibialis anterior (Schurmann et al., 2010), and mouse skinned extensor digitorum longus (EDL) muscle fibers (Forderer et al., 2016). In addition, the dynamic measurement of SHG anisotropy has been achieved in a dissected intact frog tibialis anterior muscle fiber during isometric force generation (Nucciotti et al., 2010), worm walking in *Caenorhabditis elegans* (Psilodimitrakopoulos et al., 2014) and collagenase-treated interossei cells (Forderer et al., 2016). The dependence of SHG anisotropy on longitudinal and lateral stretch stresses in cardiomyocytes was measured in a relaxed state in a chemically fixed sliced sample (Yuan et al., 2019).

**Figure 1.**
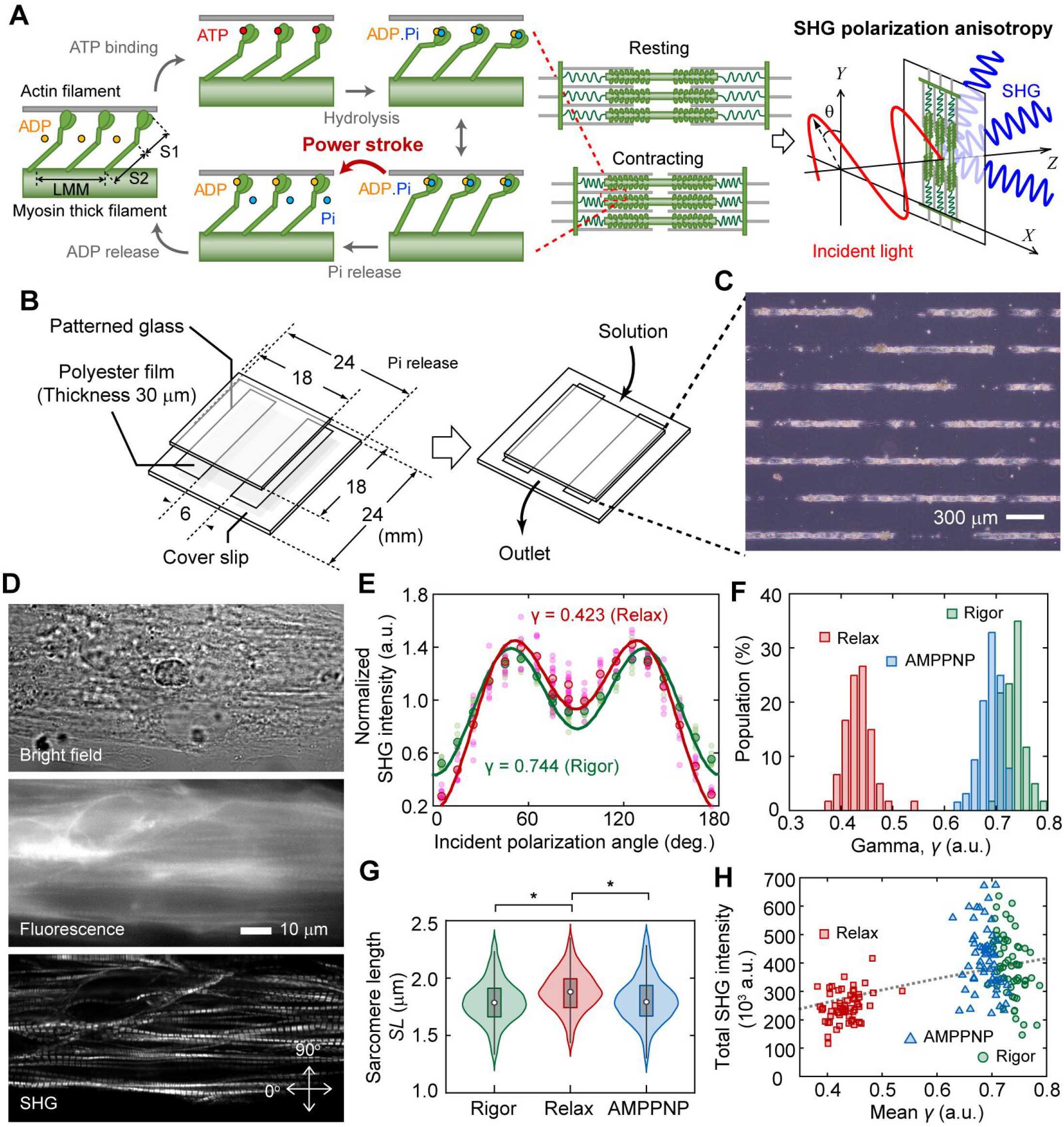
Probing structural conformation of actomyosin complex by SHG anisotropy. (**A**) Explanative drawing of concept of probing actomyosin conformation in a sarcomere by SHG anisotropy. Coupling with ATP hydrolysis, myosin changes conformation of its S1 region causing angular change of its S2 fragment, which is called mechanochemical coupling. The detected SHG anisotropy was altered by the attachment/detachment of myosin S1 to/from actin. (**B**) Explanative drawing of our sample preparation. (**C**) A microscopic photo of cells adhered on a line-and-space pattern substrate. (**D**) Typical example of SHG image acquisition of cardiomyocytes on a patterned line. Top, bright field image; middle, fluorescent image with fluorescent-phalloidin; bottom, SHG image. Arrows indicate the orthogonal axes of the incident polarization. (**E**) Incident polarization dependence of SHG emitted from a sarcomere in the absence (*green, Rigor*) or the presence of ATP (*red, Relax*). Pink circles and light green circles are typical 10 traces in the absence or the presence of ATP, respectively. Red circles and green circles are mean value of the 10 traces of them. Solid lines are fitting results with equation Eq. 2. The value of SHG intensity is normalized by the intensity of averaged value of all obtained data for relax state at 90°. (**F**) Histograms of mean value of parameter *γ* in a field of view (FOV) in the absence (*green*, N = 60 FOVs) or the presence of ATP (*red*, N = 60 FOVs), and in the presence of AMPPNP (*blue*, N = 64 FOVs). (**G**) Violin plots of sarcomere length in the absence (*green*, N = 1,339 sarcomeres) or the presence of ATP (*red*, N=1,420), and in the presence of AMPPNP (*blue*, N = 1,486). Asterisks indicate less than 0.01 of the p-value in the Student’s t-test. (**H**) Correlation between the mean *γ* and the total SHG intensity in a FOV in the absence (*green*, N = 60 FOVs) or the presence of ATP (*red*, N = 60 FOVs), and in the presence of AMPPNP (*blue*, N = 64 FOVs). Black broken line is the theoretical curve based on the integration of Eq. 2.

To the best of our knowledge, there have been no reports on measuring SHG anisotropy in beating single cardiomyocytes. In this study, we constructed an assay system to measure dynamic changes in SHG anisotropy emitted from a sarcomere in a live cardiomyocyte using a highly sensitive SHG polarization microscope with a fast polarization-controllable device, as previously described (Kaneshiro, Watanabe et al., 2016, Shima, Morikawa et al., 2018). The key gadget was a line-and-space pattern substrate, which could solve the problem of sarcomere arrangement irregularity, making the experiment difficult. Using our SHG polarization microscope and the line-and-space pattern substrate, we investigated the relationship between SHG polarization originating from actomyosin activity and disease phenotypes in two models: one based on induced pluripotent stem cell (iPSC) technology of genetic hypertrophic cardiomyopathy and another on acquired muscle failure induced by ultraviolet irradiation. Because SHG measurement is based on two-photon excitation optics, the wavelength of the irradiated light is in the infrared region, enabling deep tissue imaging. As a more challenging experiment, we further applied the present method to evaluate actomyosin dysfunction in a *Drosophila* model of Barth syndrome (Xu, Condell et al., 2006). The present actomyosin evaluation method based on the dynamic measurement of SHG anisotropy developed in this study promises to provide a new effective tool in cardiomyopathy research and medicine.

## Results

### Construction of an assay system for SHG polarization measurement in cardiomyocytes

SHG is a nonlinear scattering phenomenon that occurs in both the forward and backward directions. In general, SHG intensity is stronger during forward scattering than backward scattering (James & Campagnola, 2021), 26-fold higher in our study (Appendix Fig. S1). Forward- and backward-scattered SHGs have different implications; because the backward-scattered SHG contains signals derived from the multiple scattering of the forward-scattered SHG when observing a thick sample, the backward scattered SHG is difficult to interpret (Chu, Tai et al., 2009, Williams, Zipfel et al., 2005). Meanwhile, measuring the backscattered SHG is suitable for biological sample observations. Here, we chose a transmission SHG microscope because of its high signal strength for fast measurements during a cardiomyocyte pulsation (Appendix Fig. S2) and simple interpretation due to absence of signal contamination. In a transmission-type microscope, the sample should be placed between two objectives. A special chamber was composed of two glass plates and a polyester spacer with dimensions of 18 mm × 6 mm × 0.03 mm (Fig. 1B). A line-and-space pattern substrate allowed cardiomyocytes to attach in a linear fashion (Fig. 1C).

Here, we prepared spontaneously beating cardiomyocytes differentiated from human iPSCs (hiPSCs), 253G1 strain. The sarcomeres in chemically permeabilized cardiomyocytes were arranged along the employed line-and-space pattern and selectively visualized via SHG (Fig. 1D). The state transition of the actomyosin crossbridge from rigor to relaxation in permeabilized cardiomyocytes could be induced by loading an ATP solution into the chamber on the microscope. Following the experimental procedure in previous reports (Nucciotti et al., 2010, Plotnikov et al., 2006, Schurmann et al., 2010), we measured the SHG intensity at various incident polarizations of a sarcomere with (in relax state) or without (in rigor state) treatment with 5 mM ATP (Fig. 1E). Polarization dependence was expressed using the following equation, assuming the dipole polarity angle *φ* of the summation of the S1 and S2 directions within a confocal volume:

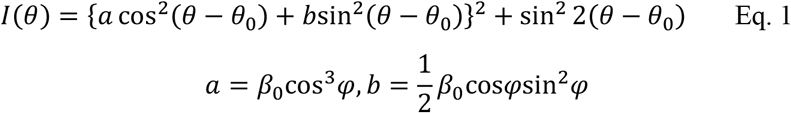

where *θ* is the incident polarization, *θ*_0_ is the orientation angle of the sarcomere in microscope coordinates, and *β*_0_ is a proportional constant (Tiaho, Recher et al., 2007). The parameter *γ*, which reflects the average crossbridge state (Nucciotti et al., 2010, Plotnikov et al., 2006, Schurmann et al., 2010), was estimated by fitting the obtained polarization dependency of the intensity using the following equation:

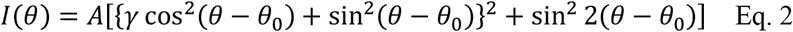

where *A* is the proportionality constant (Fig. 1E, *solid lines*).

The mean value of *γ* in a field of view (FOV) of 128 × 512 pixels (25 × 100 μm) clearly decreased from *γ*_rigor_ = 0.73 ± 0.02 (mean ± standard deviation) in rigor state, to *γ*_relax_ = 0.43 ± 0.03 in relaxed state (Fig. 1F, *red and green*). These values were consistent with the values reported in previous studies, where *γ*_rigor_ = 0.68 ± 0.01 and *γ*_relax_ = 0.46 ± 0.03 for rabbit psoas fibers (Nucciotti et al., 2010), *γ*_rigor_ = 0.73 ± 0.04 and *γ*_relax_ = 0.50 ± 0.05 for mouse tibialis anterior (Schurmann et al., 2010), and *γ*_rigor_ = 0.70 ± 0.06 and *γ*_relax_ = 0.45 ± 0.04 for mouse EDL fibers (Forderer et al., 2016). Loading 5 mM of 5’-adenylylimidodiphosphate (AMPPNP), a nonhydrolyzable analog of ATP, slightly but significantly decreased *γ* to *γ*_AMPPNP_ = 0.69 ± 0.02 (P = 5.3×10^−22^ in Student’s t-test) (Fig. 1F, *blue*). While AMPPNP was thought to induce myosin dissociation from actin, sarcomere length (*SL*) was not increased by AMPPNP treatment but was increased by ATP (P = 1.3×10^−37^ in Student’s t-test) (Fig. 1G). It was thought that the majority of myosin remained interacting with actin, even in the presence of 5 mM AMPPNP, and only a part of the dissociated myosin population contributed to the decrease in *γ* in the present case. These results strongly suggest that the dissociation of myosin from actin filaments causes a disturbance in the polarization orientation of electric dipoles in a sarcomere, coupled with a decrease in *γ*.

According to Eq. 2, the integral of the obtained SHG intensity should monotonously increase to the *γ*-value. The total SHG intensity in the FOV decreased following the decrease in *γ* after treatment with ATP, but not with AMPPNP (Fig. 1H). While a monotonous increase was observed within the ATP data (Fig. 1H, *red*), the intensity was negatively correlated with the *γ*-value within the rigor or AMPPNP data (Fig. 1H, *green and blue*), indicating the presence of a factor contributing to intensity but not related to the *γ*-value. Thus, SHG anisotropy measurement, apart from intensity measurement, could be meaningful for probing actomyosin activity.

### Dynamic measurement of SHG anisotropy in living cardiomyocytes

The SHG polarization microscope we constructed enables the measurement of the incident polarization dependence of SHG within 1 ms/pixel (Kaneshiro, Okada et al., 2019, Kaneshiro et al., 2016). By limiting the FOV to 2 × 40 pixels (0.39 × 7.81 μm), including approximately 2-3 sarcomeres, measurement within 80 ms can be possible (Fig. 2A). The dynamic spike-like change in *γ* synchronized with sarcomere contraction was successfully measured during spontaneous beating in living cardiomyocytes derived from hiPSCs (Fig. 2B and Supplementary Video S1). The average *γ* at the resting state *γ*_rest_ and the contracting state *γ*_cont_ could be estimated by fitting the histogram of *γ* with a double Gaussian distribution (Fig. 2C). The *γ*_rest_ = 0.43 obtained corresponded to the *γ*_relax_ (0.43) obtained in permeabilized cardiomyocytes, whereas the *γ*_cont_ = 0.50 was smaller than *γ*_rigor_ (0.69). This tendency was the same as that previously reported for isometric force generation in frog muscle fibers (Nucciotti et al., 2010). The value of *γ*_cont_ = 0.50 was reasonable, assuming that *γ* reflects the population of myosin bound to actin. The value of *γ*_cont_ can be predicted using the obtained *γ*_rigor_ and *γ*_relax_, (0.69 - 0.43) × 0.25 + 0.43 = 0.495, since the population of myosin contributing to force generation during contraction is only between 20–30% (Piazzesi, Reconditi et al., 2007). During the continuous beating cycle, the relationship between sarcomere length *SL* and *γ* followed approximately the same trajectory (Fig 2D), indicating a one-to-one relationship between sarcomere motion and *γ* dynamics.

**Figure 2.**
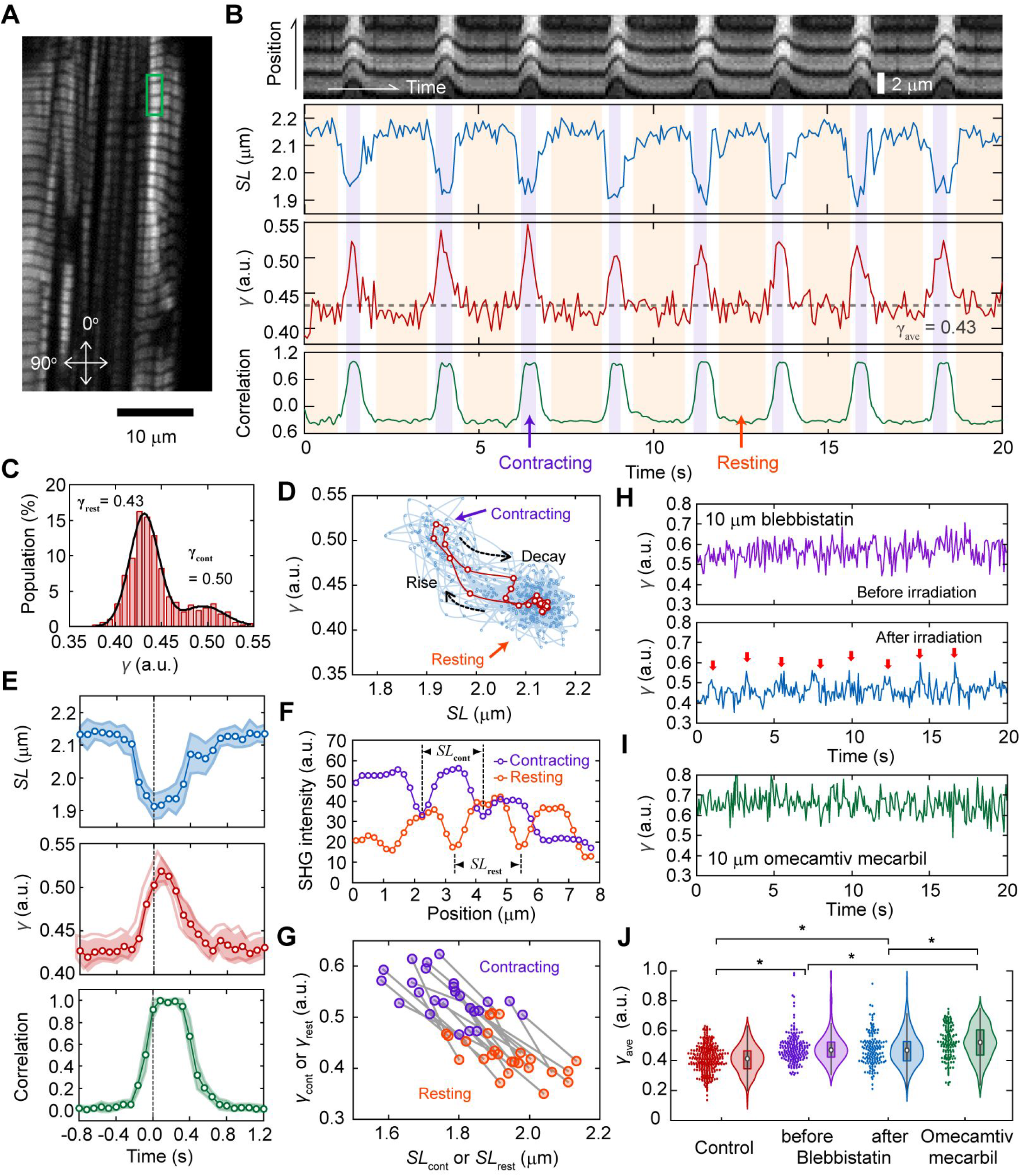
Probing dynamic structural conformation of actomyosin complex in a living cardiomyocyte. (**A**) A typical example of SHG image acquisition of a living cardiomyocytes. Arrows indicate the orthogonal axes of the incident polarization. (**B**) A typical example of dynamic measurement of parameter *γ* with 80 ms time resolution, at the area shown in green rectangle in **A**. Top, the image kymograph based on the total SHG intensity; second top, sarcomere length *SL*; second bottom, parameter *γ*; bottom, sarcomere motion converted into numerals by calculating correlation with an image at relaxation state. Light magenta and orange in the back indicates contracting and resting phases, respectively. (**C**) Histogram of *γ* within a trace of 40 sec. Black line is a fitting result with double Gaussian distributions. (**D**) Relationship between *SL* and *γ* obtained in data shown in ***B***. Cyan circles and lines are all raw data, and red ones are the average values shown in **E**, *middle*. (**E**) Average traces of *SL* (*top*), *γ* (*middle*), and sarcomere motion (correlation) (*bottom*) in 16 pulsations for 40 secs. The time zero was adjusted at the time of the largest value of *SL*. Light colors are standard deviations. (**F**) Cross-sections of SHG image averaged in contraction state (*magenta*) or resting state (*orange*). (**G**) Correlation plots between *SL*_rest_ or *SL*_cont_ and *γ*_rest_ or *γ*_cont_ of 25 traces. The pair of *SL*_rest_–*γ*_rest_ and *SL*_cont_–*γ*_cont_ obtained from a trace are linked by a gray line. (**H**) A typical trace of *γ*-value after 10 μm BS treatment (*upper*) and further after inactivation of BS efficacy by blue light irradiation (*lower*). Red arrows indicate each pulsation. (**I**) A typical trace of *γ*-value after 10 μm OM treatment. (**J**) Violin plots of *γ* in the case of no chemical treatment (*red*, N = 295 traces), 10 μm BS treatment (*magenta*, N = 176), after inactivation of 10 μm BS treatment (*blue*, N = 143), and 10 μm OM treatment (*green*, N =141). Asterisks indicate less than 0.01 of the p-value in the Student’s t-test.

Plots of the average *SL* and *γ* trace during pulsation highlighted that local sarcomere shortening preceded the increase in *γ* while the decrease in *γ* preceded sarcomere lengthening (Fig. 2E, *top and middle*), indicating that the relationship between *SL* and *γ* was nonlinear during the transition process. Average sarcomere lengths in the contracting and resting states, denoted *SL*_cont_ and *SL*_rest_, respectively, were measured using the average SHG intensity profiles during the contracting and resting states with reference to the *γ*-value (Fig. 2F and Appendix Fig. S3): the mean *SL*_rest_ and *SL*_cont_ were determined to be 1.94 ± 0.10 μm and 1.77 ± 0.10 μm (N = 25 FOVs), respectively, which were consistent with commonly known values measured in myocardial cells (Hanft et al., 2008). The correlations of *γ*–*SL* in a 40-sec time course were plotted on the same line, regardless of being the contracting or resting state (Fig. 2G). This result strongly supports the interpretation that the *γ*-value directly reflects the population of myosin interacting with actin for force generation, as previously proposed (Nucciotti et al., 2010, Schurmann et al., 2010). Instead of the *SL* measurement, the change in sarcomere shape from the relaxed state could be also quantified by the correlation between the SHG images in the resting and contracting states (Fig. 2B, *bottom*, and 2E, *bottom*). This method lost the length information and the synchronization timing between local sarcomere contraction and the *γ*-value because the motion of sarcomeres in a FOV were affected by those outside the FOV. Nevertheless, the contraction duration and relaxation delay during sarcomere motion can be obtained without care of defocusing due to muscle contraction.

By treating cardiomyocytes with 10 μM blebbistatin (BS), a widely used inhibitor of actomyosin ATP hydrolysis (Kampourakis, Zhang et al., 2018, Straight, Cheung et al., 2003), pulsation was stopped due to the inactivation of myosin force generation (Fig. 2H, *upper*). BS is known to retain the myosin ATPase cycle in the ADP.P_i_ state by inhibiting the release of phosphate (Kovacs, Toth et al., 2004). The average *γ* in a 40-sec time course, *γ*_ave_, was increased after treatment with 10 μM BS from 0.41 ± 0.09 without BS to 0.48 ± 0.10 (*p* = 4.7×10^−15^ in Student’s t-test) (Fig. 2J, *red and magenta*). By inactivating the BS effect via blue laser irradiation (Kolega, 2004, Sakamoto, Limouze et al., 2005), cardiomyocytes resumed beating, and the time course of *γ* exhibited periodic spiking, followed by a decrease in *γ* during the resting state (Fig. 2H, *lower panel*). Since not all sarcomeres showed recovery of pulsation, no significant difference was observed in the histogram of *γ*_ave_ before and after the irradiation, but a small decrease was confirmed in the mean value of *γ*_ave_, 0.47 ± 0.11 (Fig. 2J, *magenta and blue*). Treating cardiomyocytes with 10 μM omecamtiv mecarbil (OM), a cardiac-specific myosin activator and a potential pharmaceutical cure for heart dysfunction (Kampourakis et al., 2018, Malik, Hartman et al., 2011, Teerlink, 2009), remarkably reduced the occurrence of pulsation, making it difficult to capture pulsation within 40 sec of measurement (Fig. 2I). Unlike BS, which is known to interact with the nucleotide-binding pocket in S1, OM is known to bind to an allosteric site to stabilize the lever arm in a position prior to force generation, resulting in a greater population of myosin crossbridges without changing the structure of S1 (Planelles-Herrero, Hartman et al., 2017, Winkelmann, Forgacs et al., 2015, Woody et al., 2018). In view of this inhibitory mechanism, OM treatment was expected to increase *γ*_ave_ above *γ*_cont_ according to the previously proposed interpretation that *γ* reflects the crossbridge population. Indeed, the mean *γ*_ave_ was increased to 0.52 ± 0.11 (P = 1.1×10^−23^ in Student’s t-test) upon treatment with 10 μM OM (Fig. 2J, *green*). The increase in *γ*_ave_ after OM treatment was significantly larger than that by BS, both before and after inactivation (P = 0.007 and 0.001, respectively, in Student’s t-test). In summary, the dynamic measurement of SHG polarization anisotropy provided quantitative parameters to assess actomyosin crossbridge formation in living cardiomyocytes.

### Non-invasive evaluation of force generation dysfunction and repair in cardiomyocytes derived from hiPSC obtained from patients with genetic heart disease

We further elucidate the relationship between crossbridge formation and SHG anisotropy in sarcomeres as demonstration of the applicability of the present method to evaluate a dysfunctional phenotype of genetic cardiomyopathy. Mutations in the *MYBPC3* gene encoding cardiac myosin binding protein C (cMyBPC) are among the most common gene mutations found in hypertrophic cardiomyopathy (Carrier, Mearini et al., 2015, Flashman, Redwood et al., 2004). The cMyBPC function is an inhibitory control of crossbridge formation by bridging between myosin and actin. As previously reported, a deficiency in cMyBPC induces excessive myosin-actin interaction, causing an increase in contractile force generated and insufficient relaxation: a heterozygous deficiency enhances crossbridge formation, while a homozygous deficiency further inhibits super relaxation, which is the state with a 10-fold slower ATPase cycle than myosin in the conventional relaxed state (Toepfer, Wakimoto et al., 2019). Here, we established a hiPSC line with a heterozygous frameshift mutation from a patient with hypertrophic cardiomyopathy caused by an *MYBPC3* mutation (*Mybpc3*^t/+^). For comparison, hiPSC lines with a homozygous mutation (*Mybpc3*^t/t^) as a positive control or without mutations (*Mybpc3*^+/+^) as a negative control were also established with a genome editing technique. We then measured the time course of SHG polarization emitted from sarcomeres in cardiomyocytes derived from these cell lines (Fig. 3A).

**Figure 3.**
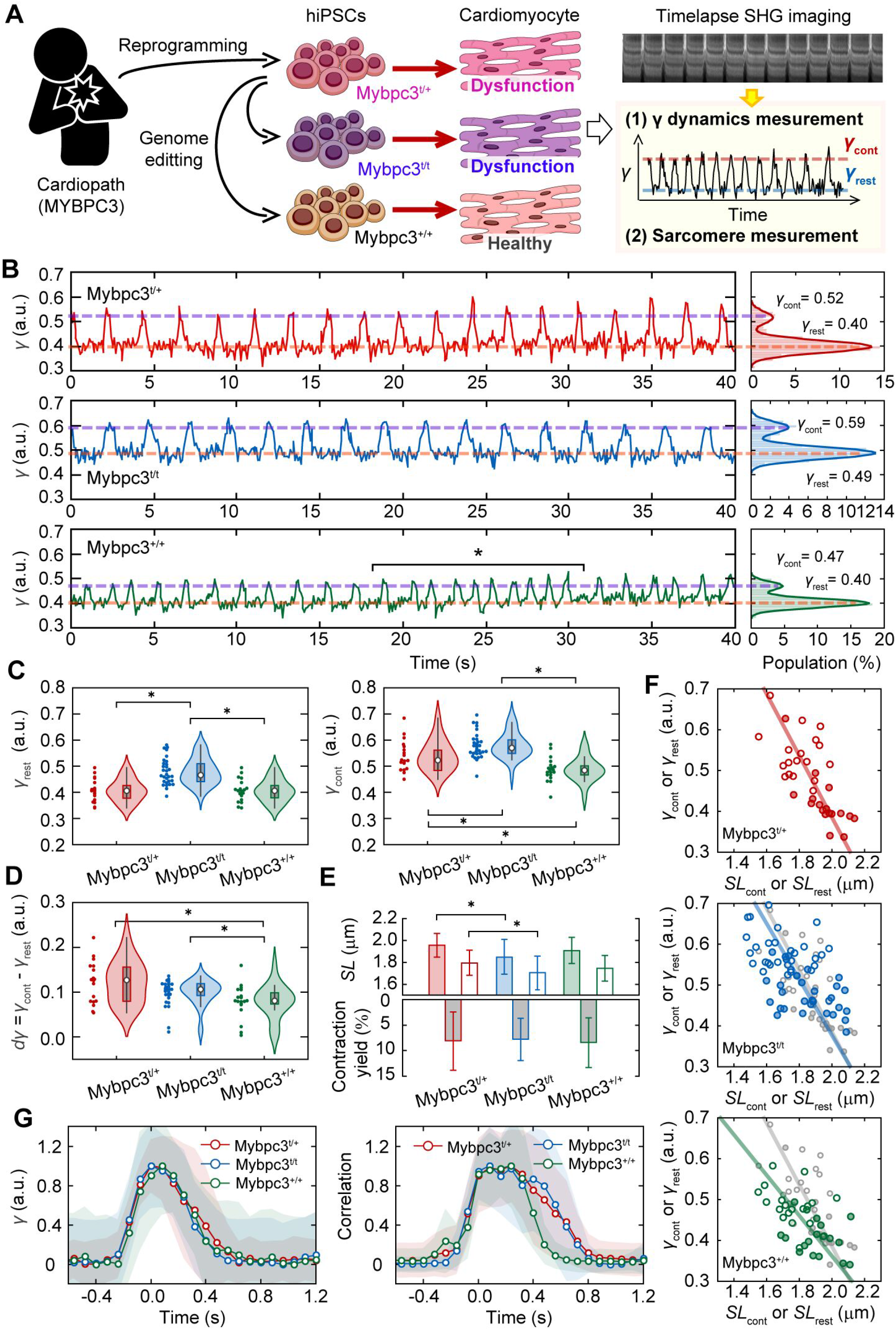
SHG anisotropy measurement of cardiomyocytes differentiated from genetic disease patient-derived hiPSCs. (**A**) A schematic drawing of the experimental procedure. (**B**) A typical example of dynamic measurement of sarcomere movement of parameter *γ* with 80 ms time resolution in *Mybpc3*^t/+^ (*top*), *Mybpc3*^t/t^ (*middle*) and *Mybpc3*^+/+^ cells (*bottom*). Right panels are histograms of *γ*-value in each trace. Solid lines in the right panels are fitting results with double Gaussian distributions. Magenta lines indicate *γ*_cont_ and orange lines indicate *γ*_rest_ in each trace obtained by the fitting. Asterisk indicates the occasional change of beating cycle. (**C**) Violin plots of *γ*_rest_ (*left*) and *γ*_cont_ (*right*) for *Mybpc3*^t/+^ (*red*, N = 21 FOVs), *Mybpc3*^t/t^ (*blue*, N = 36) and *Mybpc3*^+/+^ (*green*, N = 28). (**D**) Violin plots of *dγ*_ave_ = *γ*_cont_ - *γ*_rest_ in *Mybpc3*^t/+^ (*red*, N = 21), *Mybpc3*^t/t^ (*blue*, N = 36) and *Mybpc3*^+/+^(*green*, N = 28). (**E**) Bar graphs of *SL* in resting state (*filled*) and in contracting state (*opened*) (*upper*) and contraction yield (*lower*). Error bars are standard deviations. (**F**) Relationship between *SL*_rest_ or *SL*_cont_ and *γ*_rest_ or *γ*_cont_ for *Mybpc3*^t/+^ (*top*), *Mybpc3*^t/t^ (*middle*) and Mybpc3^+/+^ (*bottom*). Filled and opened marks are for resting state and for contracting state, respectively. Solid line is a line linked between averaged value of data for resting state and for contracting state. Overlapped gray marks and line in middle and bottom panels are those for *Mybpc3*^t/+^. (**G**) Averaged time courses of *γ*-value (*left*) and sarcomere motion (*right*) and averages trace of a pulsation in all analyzable data for *Mybpc3*^t/+^ cell (*red*, N = 8), *Mybpc3*^t/t^ cell (*blue*, N = 13) and *Mybpc3*^+/+^ cell (*green*, N = 6). Light colors are standard deviations. Asterisks in **C, D** and **E** indicate less than 0.01 of the p-value in the Student’s t-test.

Spontaneous pulsations were observed in cardiomyocytes differentiated from all established hiPSCs (Fig. 3B). We obtained the *γ*_rest_ and *γ*_cont_ values in each 40-sec time course and collected the *γ*-values from the FOVs, including a total of 60–120 sarcomeres from 17 cells. Both the mean *γ*_rest_ (0.49 ± 0.06) and mean *γ*_cont_ (0.57 ± 0.06) of *Mybpc3*^t/t^ were significantly higher than those of *Mybpc3*^t/+^ (*γ*_rest_ = 0.40 ± 0.06, P = 8.6×10^−4^; *γ*_cont_ = 0.52 ± 0.07, P = 7.6×10^−3^ in Student’s t-test) (Fig. 3C, *red and blue*). *Mybpc3*^+/+^ exhibited a lower *γ*_cont_ (0.47 ± 0.05, P = 5.7×10^−3^ in Student’s t-test) than *Mybpc3*^t/+^ while there was no significant difference in *γ*_rest_ between the two cells (0.42 ± 0.45, P = 0.73 in Student’s t-test) (Fig. 3 C, *green*). The difference between *γ*_cont_ and *γ*_rest_, denoted as *dγ* = *γ*_cont_ - *γ*_rest_ was thought to reflect the population of myosin actively interacting out of the inhibitory control. We found no significant difference between *Mybpc3*^t/+^ and *Mybpc3*^t/t^ in *dγ* (Fig. 3D, *red and blue*). Meanwhile, significant differences were observed between *Mybpc3*^+/+^ and the other cell lines (P = 3.9×10^−4^ in Student’s t-test) (Fig. 3D, *green*). Between *Mybpc3*^t/t^ and *Mybpc3*^+/+^, there were significant differences in all *γ*_cont_, *γ*_rest_, and *dγ* (P = 4.8×10^−6^, 2.2×10^−11^ and 5.4×10^−6^, respectively, in Student’s t-test). Thus, the differences in *γ* measurement represent the effect of heterozygous and homozygous cMyBPC deficiency or the repair of actomyosin activity.

While we expected that the contraction yield of sarcomeres would increase based on a previous experimental report that cellular shortening increased with cMyBPC deficiency (Toepfer et al., 2019), obvious difference was not observed in contraction yield among the three cell lines (0.08 ± 0.06 for *Mybpc3*^t/+^, 0.08 ± 0.04 for *Mybpc3*^t/t^, and 0.08 ± 0.05 for *Mybpc3*^+/+^) (Fig. 3E, *lower*). The mean *SL*_rest_ and *SL*_cont_ were respectively 1.96 ± 0.11 μm and 1.79 ± 0.11 μm for *Mybpc3*^t/+^, decreased after homozygous mutation (*SL*_rest_ = 1.85 ± 0.16 μm, P = 8.0×10^−3^; *SL*_cont_ = 1.70 ± 0.15 μm, P = 0.02 in Student’s t-test), and did not change after rescuing the phenotype (*SL*_rest_ = 1.91 ± 0.12 μm and *SL*_cont_ = 1.75 ± 0.12 μm) (Fig. 3E, *upper*). Analysis of variance (ANOVA) for the three groups exhibited no statistically significant difference in *SL* measurements (P = 0.03 for *SL*_rest_ and 0.08 for *SL*_cont_). While there might be a trend for homozygous mutations in shortening the *SL*, we were not able to show it statistically in this analysis.

Repairing this deficiency with genome editing decreased the slope of the *γ*–*SL* correlation (Fig. 3F, *top and bottom*). Since a heterozygous *MYBPC3*-deficiency increases the probability of myosin-actin interaction during contraction (Toepfer et al., 2019), the slope of the *γ*–*SL* correlation can be attributed to the probability of crossbridge formation during contraction according to this result. In addition, the myosin in heterozygous *MyBPC3*-deficiency cardiomyocytes did not fully dissociate from actin in the resting state (Toepfer et al., 2019). While the increases in *γ*_cont_ and *γ*_rest_ upon homozygous deficiency shown in Fig. 3C were confirmed by the *γ*–*SL* correlation, there was no obvious difference in the slopes of the correlations between *Mybpc3*^t/+^ and *Mybpc3*^t/t^ (Fig. 3F, *top and middle*). It can be interpreted that the *γ*-value in *Mybpc3*^t/t^ was positively offset from that in *Mybpc3*^t/+^ because of the constant residual myosin interacting with actin. Relaxation of the *γ*-value was not affected by cMyBPC deficiency regardless of its homozygosity or heterozygosity (Fig. 3G, *left*), indicating that cMyBPC is not directly responsible for the kinetics of the association-dissociation cycle of actomyosin corresponding to ATP hydrolysis. Meanwhile, the average trace of sarcomere movement exhibited a delay of relaxation in *Mybpc3*^t/+^ and *Mybpc3*^t/t^ in contrast to *Mybpc3*^+/+^ cells (Fig. 3G, *right*), which was consistent with previous results obtained in whole-cell shortening (Toepfer et al., 2019).

In summary, the *γ*-related values (Fig. 3C–D), *γ*–*SL* correlation (Fig. 3F), and the average trace of *γ* and sarcomere dynamics (Fig. 3G) represented the phenotype of muscle dysfunction in cMyBPC-deficiency cardiomyocytes, suggesting that the two parameters *SL* and *γ* could be used as parallel indicators to determine the mechanism of actomyosin dysfunction. Especially, the probability and frequency of crossbridge formation for force generation could be expressed by the slope of *γ*–*SL* correlation.

### Detection of acquired myocardial dysfunction after photodamage

Next, we investigated the feasibility of SHG polarization anisotropy to detect acquired dysfunctions in cardiac myosin activity using an experimental model of radiation exposure. Even though ionizing radiation causes the apoptosis and necrosis of embryonic stem cells (ESCs), some ESCs partially survive and maintain their trilineage differentiation potential in humans and mice (Hellweg, Shinde et al., 2020, Wilson, Sun et al., 2010). In mice, radiation-dosed ESCs could differentiate into cardiomyocytes; however, their beating was abnormal (Hellweg et al., 2020). We reproduced the acquired myocardial dysfunction using hiPSCs instead of mouse ESCs and ultraviolet (UV) irradiation instead of ionizing radiation and investigated the effect of UV irradiation on the *γ*-value and *γ*–*SL* correlation (Fig. 4A). HiPSCs were irradiated with UV light (254 nm) at energy densities of 0.00, 31.25, 62.50, and 125.00 mJ/cm^2^. While cell viability after irradiation decreased with an increase in irradiation energy to 92%, 10.5%, 4.5%, and 0.5%, respectively, more than 90% of the surviving cells positively expressed the pluripotency markers Sox2 and Oct4 (Appendix Fig. S4). These surviving irradiated iPSCs could differentiate via embryoid body (EB) formation into beating cardiomyocytes with or without UV irradiation (Supplementary Video S2).

**Figure 4.**
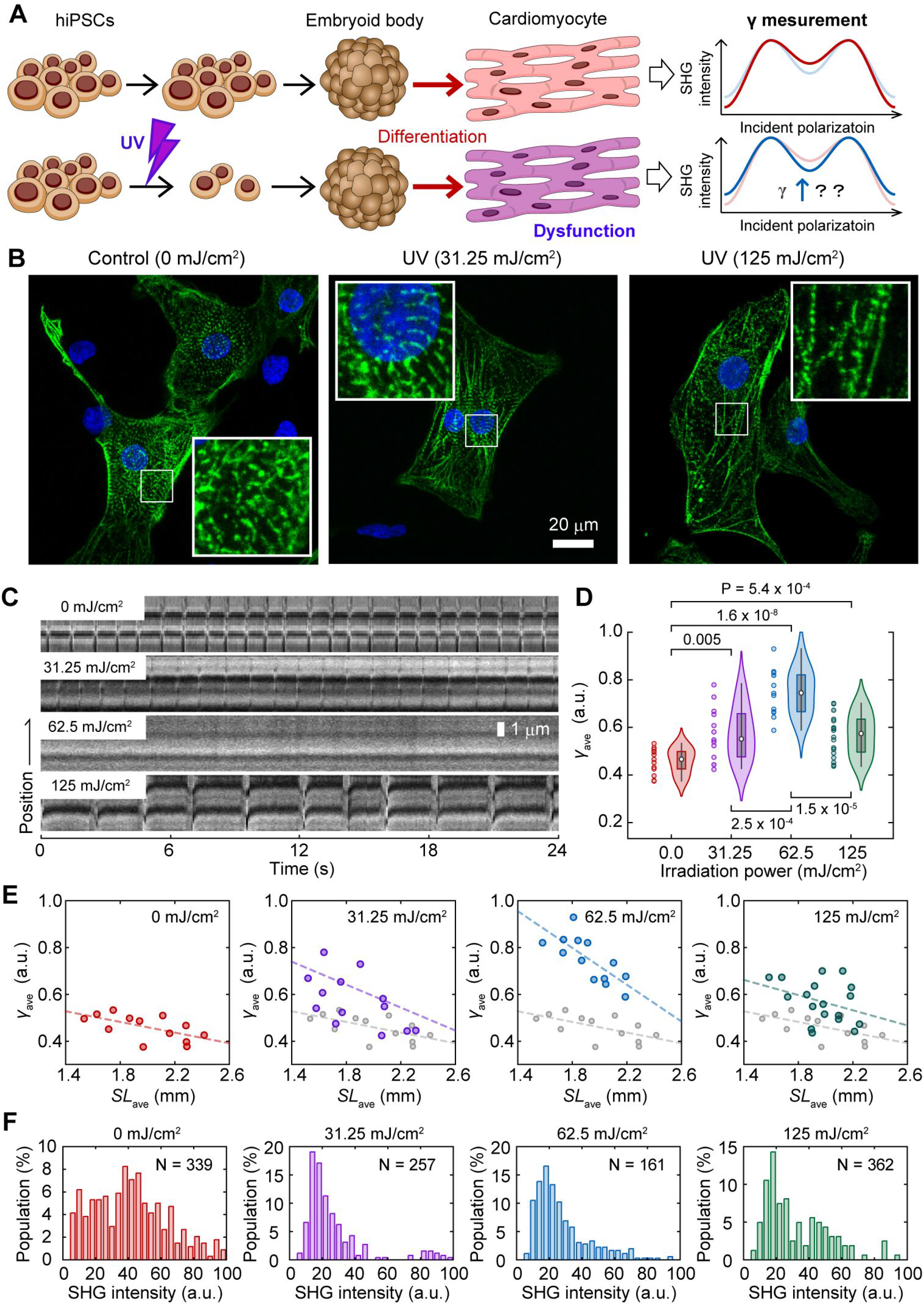
SHG anisotropy measurement of acquired force generation dysfunction. (**A**) A schematic drawing of the experimental procedure. (**B**) Confocal fluorescent microscope image of cardiomyocytes stained with anti-α-actinin (*green*) and DAPI (*blue*). Cardiomyocytes derived from hiPSCs were irradiated by UV light (254 nm) at energies of 0.00 (*left*), 31.25 (*center*), and 125.00 mJ/cm^2^ (*right*). The insets are enlarged image shown in a white rectangle in each image. (**C**) Kymographs of SHG image of sarcomeres in cardiomyocytes derived from hiPSCs irradiated at 0.00 (*top*), 31.25 (*second top*), 62.5 (*second bottom*) and 125.00 mJ/cm^2^ (*bottom*) by UV. (**D**) Violin plots of *γ*_ave_ in a sarcomere in cardiomyocytes derived from hiPSCs irradiated at 0.00 (*red*, N= 13 FOVs), 31.25 (*magenta*, N= 13), 62.5 (*blue*, N= 13) and 125.00 mJ/cm^2^ (*green*, N= 15) by UV. Asterisks indicate less than 0.01 of the p-value by the Student’s t-test. (**E**) Correlation plots between *SL*_ave_ and *γ*_ave_ for 0.00 (*red*), 31.25 (*magenta*), 62.5 (*blue*) and 125.00 mJ/cm^2^ (*green*). Solid line is a fitting result of data with a linear function. Overlapped gray marks and line in middle and bottom panels are those for 0.00 mJ/cm^2^. (**F**) Histogram of average SHG intensity of a sarcomere in cardiomyocytes derived from hiPSCs irradiated at 0.00 (*red*), 31.25 (*magenta*), 62.5 (*blue*) and 125.00 mJ/cm^2^ (*green*) by UV.

Anti-α-actinin immunostaining showed sarcomeric disarray as a sparse distribution of myocardial fibers in cardiomyocytes derived from the UV-irradiated hiPSCs, demonstrating that UV irradiation induced acquired abnormalities in myofibers during myocardial differentiation (Fig. 4B). This was also observed in mouse ESCs exposed to ionizing radiation (Hellweg et al., 2020). Even though the beating of the cardiomyocyte colonies seemed normal, the pulsations locally observed on the SHG microscope were weakened after irradiation at 31.25 mJ/cm^2^, and they became almost undetectable after UV irradiation at 62.25 mJ/cm^2^ (Fig. 4C). Surprisingly, sarcomere pulsation was still observed upon further irradiation (Fig. 4C, *bottom*). The *γ*_ave_ calculated, including both pulsating and non-pulsating sarcomeres, were 0.46 ± 0.05, 0.57 ± 0.11, 0.75 ± 0.10, and 0.57 ± 0.09 upon UV irradiation at 0.00, 31.25, 62.50, and 125.00 mJ/cm^2^, respectively (Fig. 4D).

The sarcomere length was not shortened upon UV irradiation, although the *γ*_ave_ value increased (Appendix Fig. S5). The slope of the relationship between the average *SL* (*SL*_ave_) and *γ*_ave_ changed depending on the irradiation power (Fig. 4E). UV irradiation at 31.25 mJ/cm^2^ increased both the slope and the intercept of the liner *γ*_ave_*–SL*_ave_ correlation (Fig. 4E, *second left*); the slope and the intercept were further increased at 62.5 mJ/cm^2^ irradiation (Fig. 4E, *second right*); at 125 mJ/cm^2^ irradiation, the slope returned to that for no irradiation while maintaining the increase in intercept (Fig. 4E, *right*). The increase of the intercept simply reflected the increase in myosin population that could not contribute to force generation but could still bind to actin, similar to that in the rigor state, which could presumably be inactivated or dead myosin.

According to the monotonic correlation between the total SHG intensity and *γ* shown in Eq. 2, the SHG intensity increases as *γ* increases with the irradiation power. Contrary to this expectation, the intensity of SHG emitted from the sarcomeres decreased upon UV irradiation (Fig. 4F). It can be speculated that UV irradiation decreases the number of myofibrils due to malfunction of myofibril bundling. This trial experiment demonstrated that the *γ*-value measurement allowed the evaluation of myocardial dysfunction even in cases in which sarcomere contraction rarely occurs. This emphasizes the importance of using *γ*-values rather than the SHG intensity to evaluate actomyosin activity.

### Intravital evaluation of force generation dysfunction in a *Drosophila* model of Barth syndrome

To study the effects of genetic mutations on muscle contraction and heart failure, it is desirable to conduct experiments in small animal models, such as *Drosophila*, which has a large library of gene deletions. Because one of the strong advantages of SHG microscopy is applicability to deep tissue imaging because of its two-photon excitation optics, the present method may be applicable to *Drosophila* imaging. We challenged to evaluate muscle dysfunction in a *Drosophila* model of Barth syndrome caused by mutations in the *tafazzin* gene (*TAZ*)(Pu, 2022, Xu et al., 2006). *TAZ* deficiency in humans results in cardiomyopathy, neutropenia, myopathy, growth retardation, and 3-methylglutaconic aciduria through a deficiency of the phospholipid cardiolipin (1,3-bis(sn-3’-phosphatidyl)-sn-glycerol) in the inner mitochondrial membrane (Zegallai & Hatch, 2021). In this study, we prepared a white mutant strain of *D. melanogaster, w*^1118^, as a control and a homozygous *TAZ* deficiency mutant strain (*TAZ-/-*), which exhibits Barth syndrome-related phenotypes, such as having a significant reduction in cardiolipin, mitochondrial abnormalities, and the degradation of muscle activity (Xu et al., 2006). A third instar larva from each fly strain was grown to the prepupa stage and mounted on coverslips with transparent adhesive tape to measure the SHG polarization anisotropy (Fig. 5A and B). *Drosophila* prepupae are considered technically challenging samples for optical observations because of their low transparency and the difficulty in controlling their spontaneous movement.

**Figure 5.**
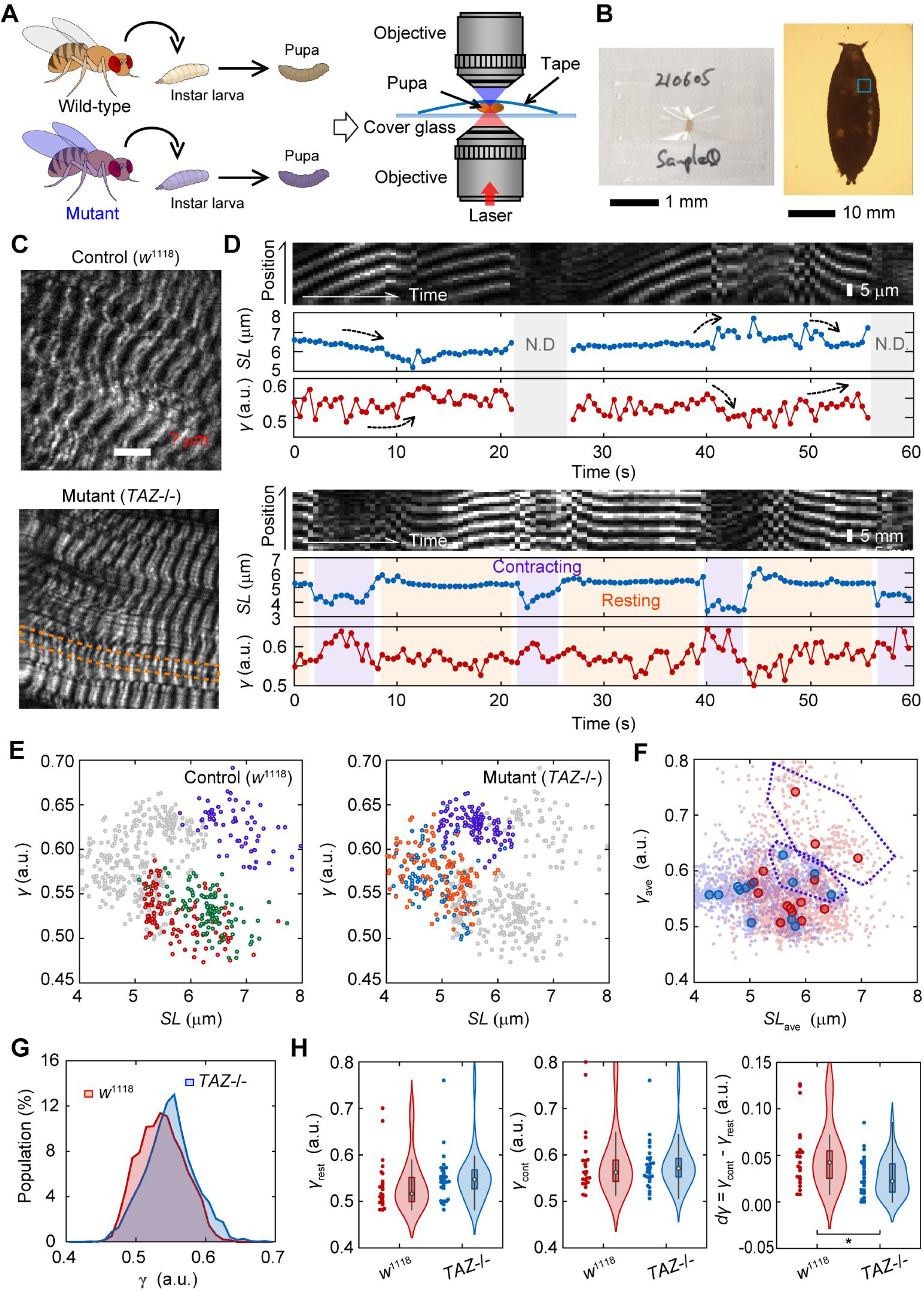
SHG anisotropy measurement in a *Drosophila* model of Barth syndrome. (**A**) A schematic drawing of the experimental procedure. (**B**) Photo graph (*left*) and micrograph (*right*) of an example of mounted *Drosophila* prepupa between a coverslip and a transparent adhesive tape. (**C**) Typical SHG image of body wall muscles in a prepupa of wild-type, control strain, *w*^1118^ (*upper*) and tafazzin deficiency, *TAZ*-/- (*lower*). (**D**) Typical two examples of image kymograph for a SHG intensity (*top*), time courses of *SL* (*middle*) and parameter *γ* (*bottom*) with 500 ms time resolution for 60 sec in a control prepupa. Gray in the back indicates the time when sarcomeres were not detectable any more. Light magenta and orange in the back indicates contracting and resting phases that could be classified, respectively. (**E**) Correlation plots between *SL* and *γ* in typical three FOVs for control (*left*) and *TAZ*-/- (*right*). Overlapped gray marks are those for the other. **(F)** Correlation plots between *SL*_ave_ and *γ*_ave_ in a FOV for w1118 (*red*) and *TAZ*-/- (*blue*). Light colors are all data for each time point. Data suspected to be outliers are surrounded by purple broken lines. (**G**) Histograms of parameter *γ* in a prepupa of control (*red*, N = 2400 time points) and *TAZ*-/- (*blue*, N = 3720) strain. The p-value of these two groups was less than 0.01 in the Mann-Whitney U-test. (**H**) Violin plots of *γ*_rest_ (*left*), *γ*_cont_ (*center*) and *dγ*_ave_ = *γ*_cont_ - *γ*_rest_ (*right*) in prepupae of control (*red*, N = 23 traces) and *TAZ*-/- (*blue*, N = 28) strain. An asterisk indicates less than 0.01 of the p-value in the Student’s t-test.

SHG microscopy visualized the sarcomeres of the body wall muscles in *Drosophila* prepupae without uncovering the puparium, regardless of the presence or absence of *TAZ* (Fig. 5C). Unlike in the cardiac muscle, the contractions in *Drosophila* prepupae did not occur periodically. Instead, spontaneous muscle activity occurs randomly everywhere at all times in a prepupa. Hence, we extended the measurement duration to 60 s with the time interval to 500 ms, and expanded the FOV to 25 μm × 20 μm with the spatial resolution of 1 μm/pixel, in order to capture the random muscle contraction. Even in a prepupa, we could confirm an anti-parallel manner of *γ*–*SL* correlation in spontaneous contraction or relaxation rather than regularity pulsations (Fig. 5D, *upper, arrows*), except during the times when sarcomeres move out of the focal plane during observation (Fig. 5D, *upper, gray*). Occasionally, clear state transitions from resting to contracting or from contracting to resting could be pursued (Fig. 5D, *lower*). The *SL* and *γ*-values within a single FOV were negatively correlated in both control and *TAZ-/-* flies (Fig. 5E), as in the case of hiPS-derived cardiomyocytes (Fig. 2D). Plotting the data obtained from 14 FOVs for the control and 13 FOVs for *TAZ-/-* with the same coordinates, the majority of data points seemed to be linearly correlated (Fig. 5F). Meanwhile, minor populations with longer *SL* showed a different correlation (Fig. 5E and F, *magenta*).

The value of *SL* (∼7 μm) for the control prepupae (Fig. 5E and F) was consistent with the previous reports (Prent, Green et al., 2008) and was obviously different from that for *TAZ-/-* prepupae (Appendix Fig. S6), which did not contradict that of a mice *TAZ*-deficiency model (Bertero, Nickel et al., 2021). Meanwhile, we could not confirm the obvious difference in *γ* time courses between them (Appendix Fig. S7), but found only a tendency of *TAZ-/-* flies to be higher than the controls as shown in the histogram values of *γ* (Fig. 5G). A slight but significant difference was confirmed in the mean values of *γ* (0.54 ± 0.03 for control; 0.55 ± 0.04 for *TAZ-/-*, P = 0.013 in Mann-Whitney U-test). Then, we extracted the traces that included obvious contracting-resting transitions and estimated the *γ*_rest_, *γ*_cont_, and the difference between them, *dγ*. Although no statistically significant difference was observed, the mean *γ*_rest_ for *TAZ-/-* flies was higher than that of the control, while the mean *γ*_cont_ remained unchanged (Fig. 5H, *left and center*). The mean *dγ* was significantly lower in *TAZ-/-* group than in the control group (Fig. 5H, *right*; P = 0.01 in Mann-Whitney U-test). Thus, the present method was applicable for investigating muscle activity in a *Drosophila* disease model.

## Discussion

In this study, we demonstrated the use of SHG anisotropy to investigate myocardial dysfunction in cardiomyopathy by constructing an effective assay system based on our highly sensitive SHG microscope with a fast polarization-controllable device (Kaneshiro et al., 2019, Kaneshiro et al., 2016), and the practical applications of this technique in examining dysfunctional cardiomyocytes differentiated from hiPSCs and a *Drosophila* disease model. Disease-derived and genome-edited cardiomyocytes provided information about the parameter *γ* obtained from SHG anisotropy measurements, reflecting myosin force generation. The sarcomere length *SL* was simultaneously measurable, and the correlation between *γ* and *SL* represented the phenotype of actomyosin dysfunction in genetically diseased cardiomyocytes. Furthermore, the present method was applicable for detecting acquired muscle failure due to radiation exposure and functional decline in *Drosophila* intravital imaging.

Two practical problems needed to be solved in measuring SHG anisotropy during myocardial beating. The first problem was the random orientation of myofibrils in cardiomyocytes, which limited the measurement of *γ*-values to well-aligned skeletal muscles. To remedy this, we used a line-and-space pattern substrate, which linearly align the myofiber orientation in cardiomyocytes (Fig. 1C and D), allowing SHG anisotropy analysis to be performed easily and reproducibly. The second problem is the low temporal resolution of current SHG polarization measurements. Cardiomyocytes beat within approximately 1 s. If the sarcomere orientation can be perfectly aligned at a given polarization angle, measurement at two or three polarization angles is sufficient to estimate the *γ*-value (Forderer et al., 2016, Psilodimitrakopoulos et al., 2014). However, unless perfectly adjusting the angle of myofiber orientation, it would be practically difficult to estimate *γ*-values under the limited angle measurement. Our highly sensitive SHG microscope enabled the measurement of SHG anisotropy at 20 angles with a 10° angle pitch within 1 ms/pixel, which provided the SHG image acquisition with a temporal interval of 80 ms including 2–3 sarcomeres during myocardial beating. One additional issue has remained to carry over to a future development, that was controlling the pulsation cycle. Thereby, we occasionally observed fluctuations in beating cycles (Fig. 3B, *bottom, asterisk*). Controlling the beating cycle needs the installation of an electrical stimulation system, buffer exchange system, and a temperature control system into a tiny space between two objectives with short working distance.

The values of parameters *γ*_relax_ (0.43) and *γ*_rigor_ (0.69) obtained in permeabilized cardiomyocytes (Fig. 1E–G) were consistent with those obtained in other specimens by other groups (Nucciotti et al., 2010, Plotnikov et al., 2006, Schurmann et al., 2010). The *γ*_rest_ (0.43) obtained in a living cell corresponded to the *γ*_relax_ in permeabilized cells, and the *γ*_cont_ (0.50) obtained was also a reasonable value, considering the active percentage (20–30%) of myosin population binding to actin during contraction (Fig. 2C–G). However, these values were far different from those previously reported during isometric tetanic contraction in single isolated fibers from frogs (*γ*_rest_ = 0.30 and *γ*_cont_= 0.64) (Nucciotti et al., 2010). This discrepancy can be explained by the difference between tetanic contraction in skeletal muscle and cardiomyocyte pulsation: more myosins interact with actin during tetanic contraction, whereas only a small portion is activated in cardiac pulsation (Brunello, Fusi et al., 2020, Hill, Brunello et al., 2021). In collagenase-treated interosseous cells, it was reported that *dγ* increased from 0.0 to 0.1 with a given electrical stimulus voltage (Forderer et al., 2016), which is in good agreement with the present results, *dγ* = *γ*_cont_ *-γ*_rest_ = 0.07. Although all the *γ*-values obtained in this study did not contradict with the values shown in the previous literatures, we think that it is not worth comparing absolute *γ*-values in living cells between different experimental conditions. As discussed in an earlier study, absolute *γ*-values largely depend on the optics and calculation theories used (Schurmann et al., 2010), and also possibly depend on species of observation sample. There should be a possibility that cell shape dependence of *γ*-value, because *γ* increases as the sarcomeres are forcibly shortened (Nucciotti et al., 2010) and that the sarcomere length depends on the surrounding environment, e.g., cell shape (Bhana, Iyer et al., 2010, Morris, Naik et al., 2020). It is more important to discuss common trends in the data obtained in the different experiments.

The attribution of *γ*, which has been controversial, is discussed here with following the present results. SHG from muscles is dominated by SHG from myosin rather than actin. Plotnikov, et al. argued that SHG from myosin is derived from the LMM and not from the S1 or S2 regions, according to their finding of small differences in SHG polarization in scallop muscles between upon AMPPNP treatment and that in the rigor state. They also did not observe any difference in SHG polarization from *C. elegans* muscles against scallop muscle, whose ratio of paramyosin (headless myosin) to myosin is 15-fold higher in *C. elegans* muscles (Plotnikov et al., 2006). Meanwhile, Nucciotti et al. and Schürmann et al. independently claimed that the main contributor to SHG is the crossbridge state, according to their observations of the differences in SHG polarization dependence between conditions where myosin attaches to actin (the rigor state) and in conditions where myosin detaches from actin (the relaxed state) (Nucciotti et al., 2010, Schurmann et al., 2010). In this study, we also confirmed the difference in *γ*-values between the rigor and relaxed states in permeabilized cardiomyocytes and between the contracting and resting states in pulsating cardiomyocytes. There is almost no doubt that SHG polarization corresponds to the crossbridge condition. It is additionally possible that the dipole orientation environment around the LMM linked to the actomyosin crossbridge condition contributes to the change in *γ*. The theoretical curve obtained using Eq. 2 almost overlapped with the averages of the data obtained in rigor state, ATP treatment, and AMPPNP treatment in the correlation plot between total intensity and *γ* (Fig. 1H, *gray broken line*). It is unlikely that the stable helical structure of LMM undergoes such a large change. This means that the primary cause of the changes in SHG polarization dependence is simply the angular change in the dipole orientation. Conclusively, it is reasonable to assume that S1 and S2 are the primary sources of change in SHG anisotropy.

Either way, the present results of the negative monotonous correlation of *γ* and *SL* (Fig. 2G), as previously reported in frog muscle fibers by Nucciotti et al. (Nucciotti et al., 2010), strongly support the previously proposed hypothesis that the *γ*-value indicates the ratio between attached to detached myosin heads (Nucciotti et al., 2010, Schurmann et al., 2010), which is a crossbridge population but not necessarily relating to force generating capability. One difference was that the *γ*_cont_ and the *γ*_rest_ depended on *SL* in cardiomyocytes, (Fig. 2G) while only *γ*_cont_ did so during isometric force generation in frog muscle fibers (Nucciotti et al., 2010). Because the *γ* obtained from myosin in the relaxed state in cardiac muscle was altered by external forces (Yuan et al., 2019), the myosin that remained undetached from actin even in the relaxed state during cardiac beating was also thought to contribute to the *γ*-value.

In the drug treatment experiments, both BS and OM, which are promising candidates as treatments for cardiomyopathy (Bond, Tumbarello et al., 2013, Roman, Verhasselt et al., 2018, Teerlink, Felker et al., 2016), treatments increased *γ*-values while stopping myocardial beating (Fig. 2H and I). Since OM treatment slows down the lever arm swinging of myosin, resulting in myosin being unable to dissociate from actin (Planelles-Herrero et al., 2017, Woody et al., 2018), the present result for OM treatment can be explained by the above hypothesis. The increase in *γ* upon OM treatment approached value close to *γ*_rigor_ (Fig 2J, *green*), which agrees with the previous result wherein OM increased Ca^2+^ sensitivity in cardiomyocytes, causing more myosin to interact with actin (Swenson, Tang et al., 2017). Meanwhile, BS treatment inhibited product release during the ATP hydrolysis cycle, resulting in myosin dissociation from actin without any interference from structural changes in the lever arm (Kovacs et al., 2004). According to the above hypothesis, BS treatment causes a *γ* decrease coupled with myosin dissociation. The actual results were contrary to this expectation and required a different or additional interpretation. It has been reported that the S1-S2 region of myosin bound to BS was more parallel to the filament axis of a myosin filament than without BS (Kampourakis et al., 2018), decreasing the dipole polarity angle *φ* in Eq. 1. In short, BS treatment is expected to cause an increase in *γ* even in non-attached myosin lying along the filament axis. A similar result was previously reported, wherein N-benzyl-p-toluene sulfonamide (BTS), another myosin inhibitor that suppresses force generation without interfering with myosin binding to actin, caused a decrease in *γ* (Schurmann et al., 2010). The *γ*-value only reflects the angle of S1 or S2 to the filament axis and is not always determined by the ratio of the populations of attached and detached myosin in specific cases, such as in chemical drug treatments. Moreover, the increase in *γ* upon BS treatment did not reach *γ*_rigor_, and the inactivation of the effect of BS treatment upon blue light irradiation restored the *γ*-value and the beating behavior (Fig 2H and J), indicating that BS only affected myosin under physiological conditions. This result does not contradict a previous report that the inhibitory effect of BS in the myocardium was not associated with calcium influx but only acted on myosin (Dou, Arlock et al., 2007). In situations wherein the mechanism of a drug’s effect on actomyosin is clarified, *γ* measurement is expected to be a powerful tool for evaluating drug efficacy.

The first demonstrative application was one of genetic cardiomyopathy, cMyBPC deficiency. We prepared cardiomyocytes differentiated from patient-derived iPSCs with heterozygous cMyBPC deficiency and those of homozygous iPSCs and repaired iPSCs produced via genome editing techniques and investigated the *γ*_cont_, *γ*_rest_, *dγ*, and *γ–SL*. An additional deficiency from the heterozygous mutant to homozygous caused an increase in *γ*_cont_ and *γ*_rest_ without changing the slope of the *γ–SL* correlation (Fig. 3C and F, *red and blue*). Rescue experiments caused a decrease in the slope of the *γ*-*SL* correlation without increasing the *γ*_rest_ (Fig. 3C and F, *green*). There are two possible causes for the increase in *γ*_rest_ according to the interpretation of the present result shown in Fig. 2; in resting state, myosin heads lying on the filament axis or the forcible interaction of myosin with actin. Considering the mechanisms underlying the effects of cMyBPC deficiency that the heterozygous mutation inhibited the super relaxation of myosin in relaxed state (Korte, McDonald et al., 2003, Toepfer et al., 2019), the latter is more plausible. The increase or decrease in the slope of the *γ–SL* correlation can be interpreted to reflect the probability of unbound myosin binding to actin during contraction. Because the slope of the *γ–SL* correlation corresponds well to the generated force, as previously reported (Nucciotti et al., 2010), this result is consistent with the finding that a cMyBPC deficiency enhances the contraction force (Toepfer et al., 2019). Interestingly, cMyBPC deficiency slowed myosin relaxation without changing *γ* dynamics (Fig. 3G), indicating a decoupling of myosin dissociation and muscle relaxation. It should be noted that the *γ*-value is only measured locally, and the sarcomere motion reflects the entire myofibril behavior in this aspect. The prolonged relaxation time mediated by the cMyBPC deficiency would not be due to myosin dissociation but rather decreased myosin cooperativity or synchrony. Although a more detailed verification is needed in the future to assess the reliability of this hypothesis, the results we obtained demonstrate that the present method allows the simultaneous evaluation of myosin dynamics both on the macro and micro scales. We therefore concluded that the present method is applicable for evaluating actomyosin dysfunction caused by cMyBPC deficiency.

Subsequently, we attempted to detect acquired myocardial dysfunction using cardiomyocyte differentiated from UV irradiated hiPSC (Fig. 4A–C). In this application, we estimated the *γ*_ave_ as an indicator to compare actomyosin activity between samples containing beating and non-beating cardiomyocytes. Curiously, the *γ*_ave_ increased depending on the irradiation power up to 62.5 mJ/cm^2^ and decreased by 125 mJ/cm^2^ irradiation (Fig. 4D). This can be explained by assuming the presence of two cell populations: cells with strong or acquired resistance to irradiation and cells damaged by irradiation. Additionally, it was also assumed that the latter population is far smaller than that of the former. At 31.25 mJ/cm^2^ irradiation, the data were mainly obtained in the damaged cell population. The damaged cells received even more damage when increasing the irradiation power to 62.5 mJ/cm^2^. A stronger irradiation dose, 125 mJ/cm^2^, killed the damaged cells, so irradiation-resistant cells were mainly detected. This interpretation is consistent with the results of the intensity histograms (Fig. 4F): the population showing high SHG intensity that disappeared at 31.25 mJ/cm^2^ irradiation reappeared with further increases in irradiation intensity. In either case that UV irradiation increased the myosin population lying along the filament axis or continuing to bind to actin without force generation, the base of the *γ* change could be attributed to the inactivated or dead myosin. Again, the increase in the slope of the *γ*_ave_–*SL* correlation is due to the promotion of crossbridge formation during muscle contraction. According to the result that both the slope and intersect of the *γ*_ave_–*SL* correlation were increased (Fig. 4E), UV irradiation might cause not only the direct inactivation of myosin function but also the indirect effect on its force generation similar to cMyBPC deficiency. The mechanism of the acquisition of dysfunction is not the focus of this study, but it is undoubtedly an interesting research prospect in radiation biology.

Finally, we applied the present method to a challenging sample: intravital imaging using a *Drosophila* disease model of *TAZ* deficiency. Although we observed a difference in *dγ* in muscle failure due to *TAZ* deficiency, some technical problems remained. First, it was difficult to select an observation area. It has been reported that regions where the sarcomeres intersect, such as those shown in Fig. 5C, *orange broken lines*, are illusory optical artifacts (Dempsey, Hodas et al., 2015). With sarcomeres constantly moving in all directions, these areas could not be avoided. Furthermore, depending on the stretching conditions and other factors, the sarcomeres may appear double-peaked or single-peaked in SHG observation (Prent et al., 2008). Thereby, the accuracy of the SHG measurement should be lower in this experiment than that in experiments using hiPSC-derived cardiomyocytes. It was also difficult to quantify the *SL* in a living *Drosophila* sample, because the sarcomeres in the field of view did not contract uniformly at the same time, and the illusory optical artifacts mentioned above also may cause false contrasts. Adopting other methods, such as using diffraction, is required to fulfill this aim. Nevertheless, we could obtain the important information to discuss the mechanism of myocardial dysfunction induced by *TAZ* deficiency. According to the previous literatures, the *TAZ* deficiency disorders mechanochemical coupling of myosin force generation via abnormal Ca^2+^ circulation induced by accumulation of reactive oxygen product (Liu, Wang et al., 2021). Meanwhile, since cardiolipin promotes ATP synthesis by stabilizing the protein complexes for the electron transport circuit on the mitochondrial membrane, the *TAZ* deficiency degrades the efficiency of ATP production by decreasing ATP synthase activity in hiPSC models (Wang, McCain et al., 2014) or by suppressing the formation of dimmer rows of ATP synthase in the *Drosophila* model (Acehan, Malhotra et al., 2011). Resulting ATP depletion would reduce myosin activity. However, Wang et al. concluded that hiPSC-derived cardiomyocyte with *TAZ* deficiency contains a critical defect in contractile force independent of ATP depletion based on their-own result that inhibition of mitochondrial ATP production in healthy cardiomyocytes has no effect on contractility (Wang et al., 2014). Summarizing the present results of a *Drosophila* Barth syndrome model, *TAZ* deficiency induced the shortening of sarcomeres and the decrease of *dγ*, most likely due to increase of *γ*_rest_, but not impacted the *γ–SL* correlation (Fig. 5). Variation of active myosin but not inactive myosin population should be expressed as a change in the slope of *γ–SL* correlation according to the previous reports (Nucciotti et al., 2010, Plotnikov et al., 2006, Schurmann et al., 2010) and the present results in Fig. 2–3. Therefore, we can claim that *TAZ* deficiency promotes the increase of inactive myosin population binding to actin, like in rigor state, but not active myosin. If ATP depletion is not involved, as Wang et al. claimed, inhibitory regulation to myosin would be promoted via disturbed Ca^2+^ flux in *TAZ* deficiency mutant. The present results in the *Drosophila* model demonstrates the applicability of the present method for deep tissue imaging is a major step forward in cardiomyopathy research.

Finally, we discuss the extensibility of SHG anisotropy measurements in actomyosin studies. Yuan et al. recently found that SHG anisotropy is attributed not only to the electric dipole along the filament axis but also to the reduction in crystallographic symmetry (Yuan et al., 2019). Crystallographic symmetry is represented as an asymmetric feature of SHG polarization. The formula for its approximate calculation was updated as follows:

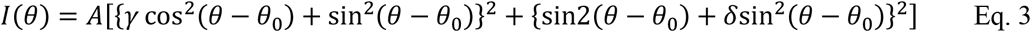

where *δ* is a parameter reflecting the crystallographic symmetry. They observed a correlation between the *δ*-value and the ratio of two myosin isoforms, α and β (Yuan et al., 2019). Asymmetric features were also observed in this study (Fig. S8A). Human atrial cardiomyocytes express two myosin isoforms, α and β, whereas ventricular cardiomyocytes express only β-myosin (Miyata, Minobe et al., 2000). In the current cardiomyocyte differentiation protocols, pacemakers, atria, and ventricles are usually generated from the same stem cells *in vitro* (Yechikov, Copaciu et al., 2016). Because the present study did not perform subtype segregation, we might have unintentionally evaluated atrial cardiomyocytes that exhibited asymmetric features. Consequently, the distribution of the obtained *δ*-values was slightly positively biased (Fig. S8B) and there were significant differences in *δ* among the rigor, relaxed, and AMPPNP-treated states (Fig. S8C). Because the strong interaction of myosin with actin filaments loads rotational tension onto myosin filaments, altering the crystallographic symmetry (Huxley, Simmons et al., 1983), the decrease in *δ* upon the addition of ATP or AMPPNP was speculated to be due to the release of rotational tension resulting from the dissociation of myosin from actin. Although the attribution of *δ*-values remains controversial, there is no doubt that *δ* measurements, along with *γ* measurements, are useful for quantifying the muscle condition.

In conclusion, this study demonstrated the applicability of SHG anisotropy to evaluate the phenotypes of genetic or acquired cardiomyopathy and quantitatively investigate its impact on actomyosin activity in cell models and a *Drosophila* model using the parameters *γ* and *SL*. Although SHG polarization microscopy cannot directly measure the actual force exerted by the muscle, the *dγ*-value reflects population of active crossbridges and the slope of *γ–SL* correlation denoting *dγ* per sarcomere contraction distance can be a measure of the relative force. The proposed method is applicable to various samples. In particular, it is expected to be useful and effective in iPS research. The cell-based disease models based on hiPSC technology promise to accelerate understanding of pathogenic mechanisms, the development of regenerative medicine, drug toxicity screening, and drug discovery (Bellin, Marchetto et al., 2012, Grskovic, Javaherian et al., 2011). To evaluate drug toxicity, disease severity, or the efficacy of a drug against hiPSC-derived cardiomyocytes, measurements of myocardial activity, e.g., muscle force generation, are required (Miyagawa & Sawa, 2018, Sewanan & Campbell, 2020). Considering the general use of hiPSCs, it is desirable to use quality-checked cells directly for this purpose. SHG microscopy, which is a non-staining and non-invasive method, has an advantage in this regard. In fact, a technique has been proposed to identify subtypes of hiPSC-derived cardiomyocytes by non-invasively measuring the total SHG intensity of each cell, even without staining (Chang, Kao et al., 2020). In addition, the use of *δ* measurements may also improve the precision of the subtype identification. This advantage is now expanding its application in the diagnosis of cancers, fibrosis, diseases involving the cornea, and tissue engineering (James & Campagnola, 2021). We hope that the present method will be an essential tool in mechanobiology, cardiomyopathy, and iPS research in the future.

## Materials and Methods

### iPSC culture and cardiomyocyte differentiation

The 253G1 human iPSC line was purchased from the Riken Cell Bank (HPS0002) and was adapted to feeder-free conditions as described previously (Nakagawa, Koyanagi et al., 2008). The *Mybpc3*^t/+^ hiPSC strain was established from patient, and the *Mybpc3*^t/t^ and *Mybpc3*^+/+^ strains were edited using CRISPR-Cas9, as previously reported (Takeda, Miyagawa et al., 2020, Higo, Hikoso, et al., 2021). The cells were maintained on iMatrix511-silk-coated (892021, Matrixome) for 253G1 or iMatrix-511-corted (892012, Matrixome) dishes for *Mybpc3*^t/+^, *Mybpc3*^t/t^, and *Mybpc3*^+/+^ strains in StemFit AK02N medium (Ajinomoto) at 37°C and 5% CO_2_. The medium was exchanged every day, and passaging was performed twice per week.

For cardiomyocyte differentiation, hiPSCs were harvested with TrypLE select (12563011, Gibco), transferred to ultra-low attachment U-bottom 96 well plate (MS-9096U, Sumitomo Bakelite) at a density of 1 × 10^4^ cells/well, and then acclimated to the mTeSR1 culture medium (ST-85850, STEMCELL Technologies) until differentiation began. Differentiation was induced 4 days after EB formation using the PSC Cardiomyocyte Differentiation Kit (A2921201, Gibco), following the differentiation time course described in the manufacturer’s instructions. On 2 days after the induction, EBs were transferred from 96 well plate to gelatin/iMatrix/fibronectin triple-coated dish (concentrations: 0.1%, 0.5 µg/cm^2^ and 1 µg/cm^2^, respectively) and allowed to differentiate up to 14 days after the induction.

### UV irradiation

HiPS cells on culture dishes were irradiated with 254 nm ultraviolet light (UV) for 0, 2.5, 5.0, or 10 min. The UV dose was measured with an S425C thermal power sensor equipped on a PM100D monitor (Thorlabs Japan), and was calculated to be 12.5 mJ/cm^2^/min. Under these conditions, approximately 92%, 10.5%, 4.5%, and 0.5% cells survived, respectively (n = 2). These cells were cultured for more than two weeks after UV irradiation and used for subsequent experiments. We confirmed that more than 90% cells remained to express both pluripotency markers Sox2 and Oct4 markers after 10 passages with immunostaining analysis (93, 97, 95, and 99% for Sox2, and 99, 99, 100 and 99% for Oct4, in 300, 67, 351, and 100 cells, respectively) (Appendix Fig. S4).

### Immunocytochemistry

Immunostaining was performed 14 days after the induction of differentiation. Cultured cells were washed twice with phosphate-buffered saline (PBS(-)), dissociated by trypsinization (0.25% Trypsin/EDTA, 25200056, ThermoFisher Scientific), and re-plated on iMatrix511-coated cover glass (Matsunami, C218181, Japan) in culture medium consisting of Dulbecco’s modified Eagle’s medium (FujiFilm Wako Pure Chemicals) with 10 % fetal bovine serum (DMEM/10%FBS). Four days after the plating, cells were fixed with 4% paraformaldehyde (PFA) for 15 min at RT. After washing, cells were permeabilized with 0.3% Triton X-100 in PBS and then incubated with anti-α-actinin mouse monoclonal antibody (1:100, A7732, Sigma-Aldrich) in CAS-Block reagent (Life Technologies, 008120) at 4°C, overnight. After washing, immunoreactive cells were determined using the appropriate fluorescently labeled secondary antibodies, Alexa Fluor 488 conjugated goat anti-mouse IgG (1:500, Molecular probe, A11001). The cells were washed, stained with DAPI (1 μg/mL, Sigma, D9542), and mounted in Vectashield mounting medium (Vector Laboratories, H-1000).

### Seeding cardiomyocytes on a micropatterned substrate

A strong dependence of myofibril orientation in cardiomyocytes on cell shape has been previously reported (Bhana et al., 2010, Morris et al., 2020). Cell shape can be easily controlled by culturing cells on a substrate on which adhesion areas are biochemically controlled (Bray, Sheehy et al., 2008). In order to make approximately one-dimensional myofibril arrays along the line direction, we cultured all cardiomyocytes on a line-and-space patterned glass substrate with a line width of 20 μm and a space between adjacent lines of 300 μm (CytoGraph L20S300, Dai Nippon Printing Co. Ltd.) (Fig. 1C).

For seeding of 253G1-derived cardiomyocytes, the line-and-space patterned glass substrates were placed in 6 well cell culture plate (3516, Corning, NY) with the cell adherent side up, coated with iMatrix 511-silk (0.5 µg/cm^2^) in PBS(-) at 4°C overnight, and then washed three times with PBS(-). For *Mybpc3*^t/+^, *Mybpc3*^t/t^, and *Mybpc3*^+/+^-derived cardiomyocytes, the patterned glass substrates were coated with 0.1 % gelatin instead of iMatrix 511-silk in PBS(-) at 37°C for more than 30 min, and the excess gelatin solution was aspirated before seeding. On 14 day after the differentiation induction, the cardiomyocytes differentiated from hiPSCs were dissociated with 0.25% trypsin/EDTA for 10 min and resuspended in DMEM/10%FBS. Large cell aggregates were removed using a cell strainer (100 µm, 352360, Corning, NY). The remaining cells were seeded on an iMatrix-coated line-and-space patterned glass substrate at a density of 4 × 10^4^ cells/cm^2^. The culture medium was exchanged every other day until 18 days after the induction, when the SHG analysis was performed.

### Preparation of permeabilized cardiomyocytes

To investigate the nucleotide dependence of actomyosin structure, permeabilized cardiomyocytes were prepared following a previously reported protocol (Sato, Jung et al., 2017). Briefly, surface membrane of cardiomyocytes cultured on a patterned substrate was removed with extraction buffer (30 mM imidazole, pH 7.5, 70 mM KCl, 1 mM EGTA, 2 mM MgCl_2_, 0.5% Triton X-100, and 4% polyethylene glycol (mol wt 8,000)) for 4 min on ice, then washed twice with PBS. The permeabilized cells were then treated on ice with the reported solutions to induce rigor and relaxation (5 mM ATP) states (Nucciotti et al., 2010). The AMPPNP state was induced using the same solution as that used for rigor with 5 mM AMPPNP.

### Drug treatment of cardiomyocytes

(S)-(-)-Bbbistatin (BS) was purchased from Toronto Research Chemicals Inc. (TRC) (B592500, lot no. 5-FRU-48-2), dissolved in dimethyl sulfoxide (DMSO) to prepare a 10 mM stock solution, and stored at −80°C. Omecamtiv mecarbil (OM) was purchased from TRC (C544000, lot no. 1-TIM-105-1), dissolved in DMSO to make a 10 mM stock solution, and stored at −80°C. Cardiomyocytes differentiated from 253G1 hiPSCs through EB formation were plated on a patterned substrate coated with iMatrix, as mentioned above, and were incubated at 37°C in the DMEM/10%FBS with 10 μM of BS or 10 μM of OM with 0.1% DMSO for 60 min. As control sample against drug-treated cells, cardiomyocytes differentiated from 253G1 hiPSCs were incubated at 37°C with 0.1 % DMSO for 60 min. All biological assays were repeated at least thrice.

### Preparation of *Drosophila* prepupa sample

The *TAZ*-/-strain was established following the previous report (Xu et al., 2006). A chromosome with null *TAZ* mutation was balanced over the CyO (Curly of Oster) chromosome tagged with green fluorescent protein (GFP). The pupae were raised at 25 °C until the 3^rd^ instar larval stage. GFP-negative larvae were counter-selected as homozygous flies for *TAZ*. SHG images were acquired during the prepupal stage at a controlled room temperature of 23 °C. A prepupa was placed on a glass slip such that its dorsal surface was in contact with the glass surface. The sample was then immobilized from the ventral surface using transparent adhesive tape (Fig. 5B).

### Optical setup of SHG microscope

We used the same optical setup of the SHG microscope as that used in previous studies (Kaneshiro et al., 2019, Kaneshiro et al., 2016). The brief descriptions are provided below. The laser source was a Ti: sapphire pulsed laser with a wavelength of 810 nm, pulse width of 200 fs, and a repetition rate of 80 MHz. The polarization state of the incident beam was controlled using a polarization controller consisting of a pair of electro-optic modulators. The incident beam was focused by an air-immersion objective lens with a magnification of 40× and a numerical aperture (NA) of 0.95. The incident beam was raster-scanned in the observation plane by *x*-*y* galvanometer mirrors. Forward-scattered photons were detected by a photon-counting photomultiplier tube after collection by an objective lens (with a magnification of 60× and NA of 1.42 for cardiomyocytes, and with a magnification of 60× and NA of 0.70 for *Drosophila* prepupae), passed through a tube lens and a band-pass filter.

### Image data acquisition

A set of polarization-resolved SHG (p-SHG) images was acquired by synchronous control of the *x*-*y* scanner and the polarization controller. For all the measurements, the average power of the incident laser was set to 26 mW. This value led to an approximate average illumination intensity of 3 MW/cm^2^ at the focus. For cardiomyocytes, the laser focus was raster-scanned in a rectangle of 25 × 50 μm^2^ with 128 × 256 pixels, with a dwell time per pixel of 1 ms. The incident polarization was swept within the dwell time at every pixel position. The angle pitch was 10° and the number of angles was 20, which covered a total angle range of over 180°. The total acquisition time per image was about 33 s. In the dynamic measurements on 253G1 and *MYBPC3* cardiomyocytes, 500 polarization-resolved image sets within a rectangular region of interest of 0.39 × 7.81 μm^2^ (2 × 40 pixels) were repeatedly acquired with a pixel dwell time of 1 ms and no interval time between images. In the dynamic measurement of UV-treated cardiomyocytes, the rectangular region was set to 0.195 × 4.88 μm^2^ (2 × 50 pixels). The total acquisition time per 500 frames was 40 sec. For the dynamic measurement in a *Drosophila* prepupa, the rectangular region was set to 25 × 20 μm^2^ with the spatial resolution of 1 μm/pixel and the temporal resolution of 1 ms. The polarization sweeping condition was the same as that used for cardiomyocytes. A total of 120 polarization-resolved image sets were acquired within 60 s.

### Estimation of whole sarcomere motion with correlation calculation

In some experiments, *SL* measurement was not possible because of the deterioration of the SHG image during contraction. To estimate sarcomere motion in such cases, we utilized the correlation between images because SHG images in the resting state are similar, whereas those during contraction differ from resting SHG images. The correlation was calculated by first calculating and averaging Pearson’s correlation between the given time t and the others. Then, resting time was identified using Otsu’s binarization method, and Pearson’s correlation was calculated and averaged for images at a given time t and resting time images to give a correlation value.

## Acknowledgements

We thank Kohei Kawaguchi (RIKEN) for experimental preparations and research members at Dai Nippon Printing Co., Ltd. for providing the line-and-space pattern substrates and advice on how to use them. The *TAZ* mutant strain was kindly gifted by Dr. Mindong Ren (Xu et al., 2006). This work was mainly supported by the Japan Agency for Medical Research and Development (grant number: 17bm0804008 to T. M. W. and S. M.) and the Japan Science and Technology Agency (CREST: JPMJCR1852 to T. M. W. and E. K.), and was partially supported by MEXT Grant-in-Aid for Scientific Research on Innovative Areas “Singularity Biology” (grant number JP18H05409). We would like to thank Editage (www.editage.com) for the English language editing.

## Ethics statement

Research plan related to the hiPSCs used in this study were approved by the Ethics Committee of RIKEN Center for Biosystems Dynamics Research (approval number: RIKEN-K1-2021-003). The use of patient-derived samples and genomic analysis were approved by the Ethics Committee of Osaka University Hospital, and written informed consent was obtained from all patients. This study conforms to the ethical guidelines for medical and health research involving human participants in Japan and all principles outlined by the Declaration of Helsinki.

## Author contributions

T. M. W. lead this study, analyzed the data, and wrote the manuscript. J. K. constructed the microscope system and performed experiments and analyses. M. T., S. H., T. K., Y. A., Y. S. and S. M. provided cardiomyocytes from the disease-derived hiPSCs and established the protocol of the cardiomyocyte microscope observations. R. Y. and K. S. investigated the acquired dysfunction by UV irradiation. D. U. and E. K. contributed to *Drosophila* experiments. H. F. wrote the manuscript and provided a critical discussion.

---

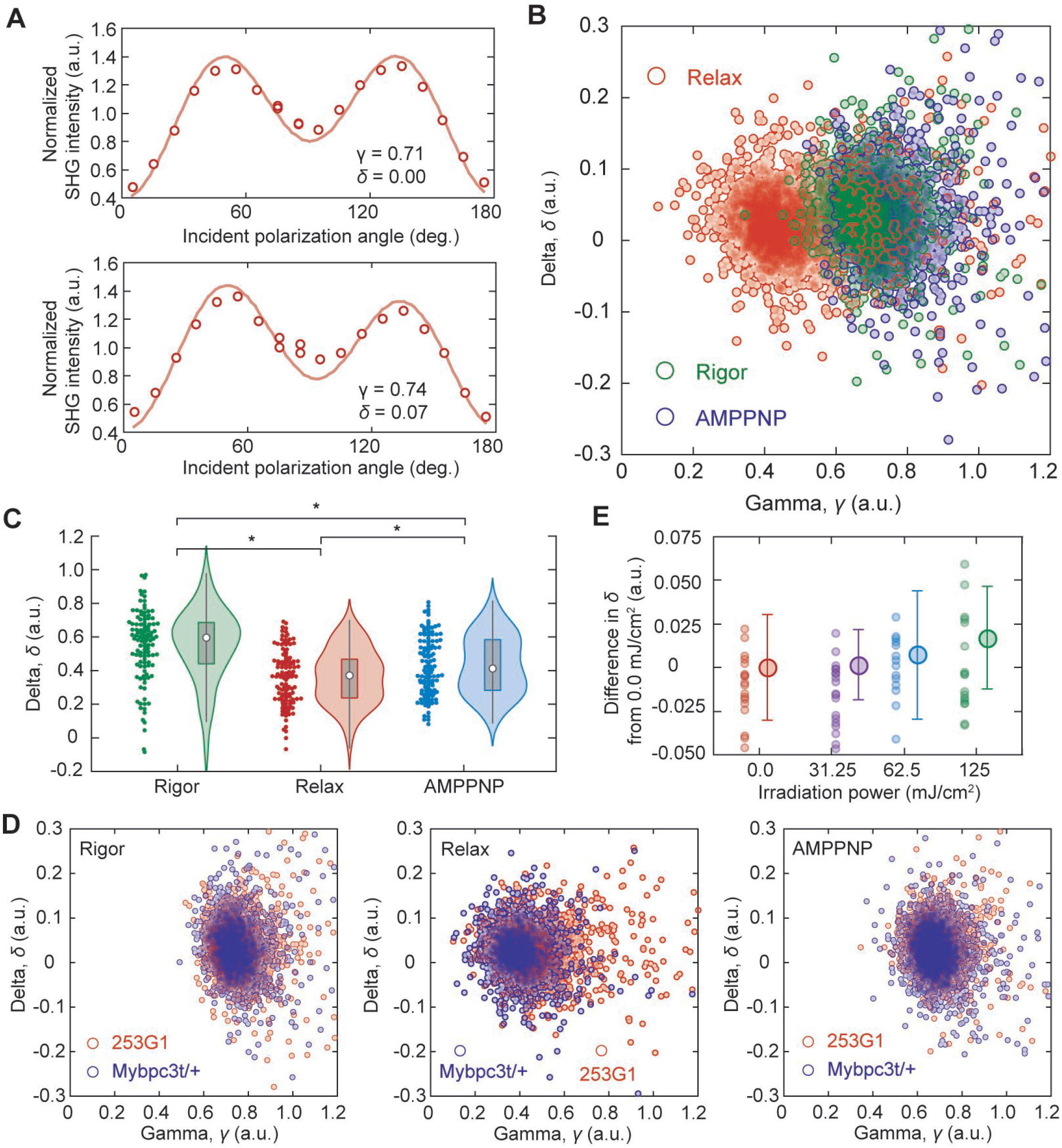

